# Neuropeptide signalling shapes feeding and reproductive behaviours in male *C. elegans*

**DOI:** 10.1101/2021.11.29.470485

**Authors:** Matthew J. Gadenne, Iris Hardege, Djordji Suleski, Paris Jaggers, Isabel Beets, William R Schafer, Yee Lian Chew

**Affiliations:** Molecular Horizons, University of Wollongong and Illawarra Health and Medical Research Institute, Wollongong Australia; Neurobiology Division, MRC Laboratory of Molecular Biology, CB2 0QH Cambridge United Kingdom; Department of Biology, KU Leuven, 3000, Leuven, Belgium; Flinders Health and Medical Research Institute and College of Medicine and Public Health, Flinders University, Adelaide, 5042, South Australia, Australia

**Author notes:** these authors contributed equally to this work (Matthew J. Gadenne, Iris Hardege).

**Keywords:** *Caenorhabditis elegans*, feeding, neuropeptides, reproduction, sexual dimorphism, sex-specific gene expression

## Abstract

Sexual dimorphism occurs where different sexes of the same species display differences in characteristics not limited to reproduction. For the nematode *Caenorhabditis elegans*, in which the complete neuroanatomy has been solved for both hermaphrodites and males, sexually dimorphic features have been observed both in terms of the number of neurons and in synaptic connectivity. In addition, male behaviours, such as food-leaving to prioritise searching for mates, have been attributed to neuropeptides released from sex-shared or sex-specific neurons.

In this study, we show that the *lury-1* neuropeptide gene shows a sexually dimorphic expression pattern; being expressed in pharyngeal neurons in both sexes but displaying additional expression in tail neurons only in the male. We also show that *lury-1* mutant animals show sex differences in feeding behaviours, with pharyngeal pumping elevated in hermaphrodites but reduced in males. LURY-1 also modulates male mating efficiency, influencing motor events during contact with a hermaphrodite. Our findings indicate sex-specific roles of this peptide in feeding and reproduction in *C. elegans*, providing further insight into neuromodulatory control of sexually dimorphic behaviours.

## Introduction

Sexual dimorphism is where two sexes of the same species show differences in behaviour or anatomical features not limited to reproduction, such as body size or muscle mass. These sex differences are important to promote organismal reproductive fitness and survival (Fairbairn, 1997). In the mammalian nervous system, sexual dimorphisms arise due to a combination of sex differences in the genome, hormonal influence, and differences in signalling within and between neural circuits (McCarthy & Arnold, 2011). It is not yet well-understood precisely how sexual dimorphism arises in the brain, or how these contribute to differences in behaviour or in the incidence of some neurological conditions (Clayton, 2016). Increasing evidence suggests that neuromodulator signalling contributes to these sex differences (see for example (Asahina *et al*, 2014; Liu *et al*, 2007; Tabatadze *et al*, 2015)); however, the size and complexity of the mammalian nervous system means that there is a relatively limited mechanistic understanding of neuromodulator networks, and how they interact with neural circuits, in these organisms. Investigating the specific effects of neuromodulators on sex differences in the brain and on behaviour in a more compact and tractable system may help clarify the general principles through which these internal signalling mechanisms regulate sexual dimorphism.

The two sexes of *Caenorhabditis elegans* are anatomically and behaviourally distinct. Hermaphrodite gonads produce both sperm and eggs, allowing them to self-fertilise and reproduce independently of male worms. Hermaphrodites do not mate with other hermaphrodites. In contrast, as males only produce sperm, male reproduction is dependent on mating with a hermaphrodite. Male worms also have more neurons, including 91 male-specific neurons, many of which are involved in coordinating reproductive behaviours (Barr *et al*, 2018). In *C. elegans*, neuroanatomical differences between the sexes arise from a combination of sex-specific programmed cell death, differentiation, and neurogenesis (reviewed in (Barr *et al.*, 2018; Portman, 2017)).

In addition to sexually dimorphic features in neural circuits, including differences in synaptic connectivity (Bayer *et al*, 2020; Cook *et al*, 2019), neuromodulators can act differently in the two *C. elegans* sexes to drive sex-specific behaviours (Barrios *et al*, 2012; Reilly *et al*, 2021). One such class of neuromodulators, called neuropeptides, have been shown in multiple experimental models to be required for broad modulatory actions (Bargmann, 2012), due to their ability to trigger responses “extrasynaptically” between neurons not physically connected by synapses (Bentley *et al*, 2016). *C. elegans* provides an excellent system to study how neuropeptides control sex-specific behaviours: it is genetically tractable, amenable to a diverse technological toolbox for interrogation of the nervous system, and has a largely complete neuronal connectome in both sexes (Cook *et al.*, 2019; Walker *et al*, 2017; White *et al*, 1986; Witvliet *et al*, 2021). Understanding how neuropeptides modulate sexually dimorphic behaviours in *C. elegans* may provide a platform to investigate how similar modulatory systems function in bigger brains.

In animals that reproduce sexually, including *C. elegans*, two key appetitive behaviours are feeding/food-seeking and mate-seeking. Male *C. elegans* experience both a reproductive pressure of having to find mates to pass on genetic traits, as well as a competing ‘feeding’ selective pressure that drives them to seek abundant food (Ryan *et al*, 2014; Wexler *et al*, 2020). A characteristic behaviour of male worms is that in the presence of food but absence of mates, males leave food to prioritise mate searching (Barrios, 2014; Lipton *et al*, 2004). This behaviour is regulated by neuropeptide signalling from both sex-shared and sex-specific neural circuits (Barrios *et al.*, 2012; Garrison *et al*, 2012).

Here, we investigated the role of a specific neuropeptide, encoded by the gene *lury-1* (luqin RYamide), in modulating mating and food-seeking behaviours in *C. elegans* males. A previous study of LURY-1 peptides in *C. elegans* focused on phenotypes in hermaphrodite animals, showing that these peptides inhibit feeding and egg-laying in a food-dependent manner through interactions with the neuropeptide receptor NPR-22 (Ohno *et al*, 2017). In hermaphrodites, these LURY-1-dependent behaviours are triggered by peptide release from pharyngeal M1 and M2 neurons. In this study, we show that *lury-1* has a different expression pattern and impact on feeding behaviour in the two sexes. In addition, LURY-1 signalling also modulates mating behaviour in male worms. Our results identify for the first time a sexually dimorphic role of LURY-1 neuropeptides in *C. elegans*.

## Results and Discussion

When investigating the expression pattern of *lury-1* using a polycistronic fluorescence reporter line (*Plury-1(3.4kb)∷lury-1 gDNA + UTR∷SL2-mKate2*), we discovered additional cells expressing this reporter in male *C. elegans* compared with hermaphrodites (**Figure 1A, 1B**). As previously reported, expression in hermaphrodites is limited to pharyngeal neurons, identified as M1 and M2 neurons (Ohno *et al.*, 2017) (**Figure 1A**). In males, expression of *lury-1* is observed in the pharyngeal neurons, as well as in cells of the male tail (**Figure 1A**), including at least one ray neuron. No expression in the hermaphrodite tail was observed in our study (**Figure 1A**) or in previous reports. As first demonstrated in (Ohno *et al.*, 2017), we independently found that LURY-1 neuropeptides can activate the G protein-coupled receptor NPR-22 with high potency (dose response curve and EC50 values indicated in **Figure 1C**) using an *in vitro* screening protocol (Beets *et al*, 2012). The expression pattern of NPR-22 in hermaphrodites was previously reported in (Palamiuc *et al*, 2017; Turek *et al*, 2016) and includes expression in the pharynx, head neurons, and intestine. Using a GFP-tagged fosmid reporter line for NPR-22, we found that expression in male *C. elegans* is observed in the pharynx, and additionally in cells of the male tail including hypodermal support cells and potentially the hook structure (**Figure 1B**). These data suggest that LURY-1 could signal to NPR-22 in the pharynx to modulate feeding, and to NPR-22 in male tail structures to modulate copulation.

**Figure 1:**
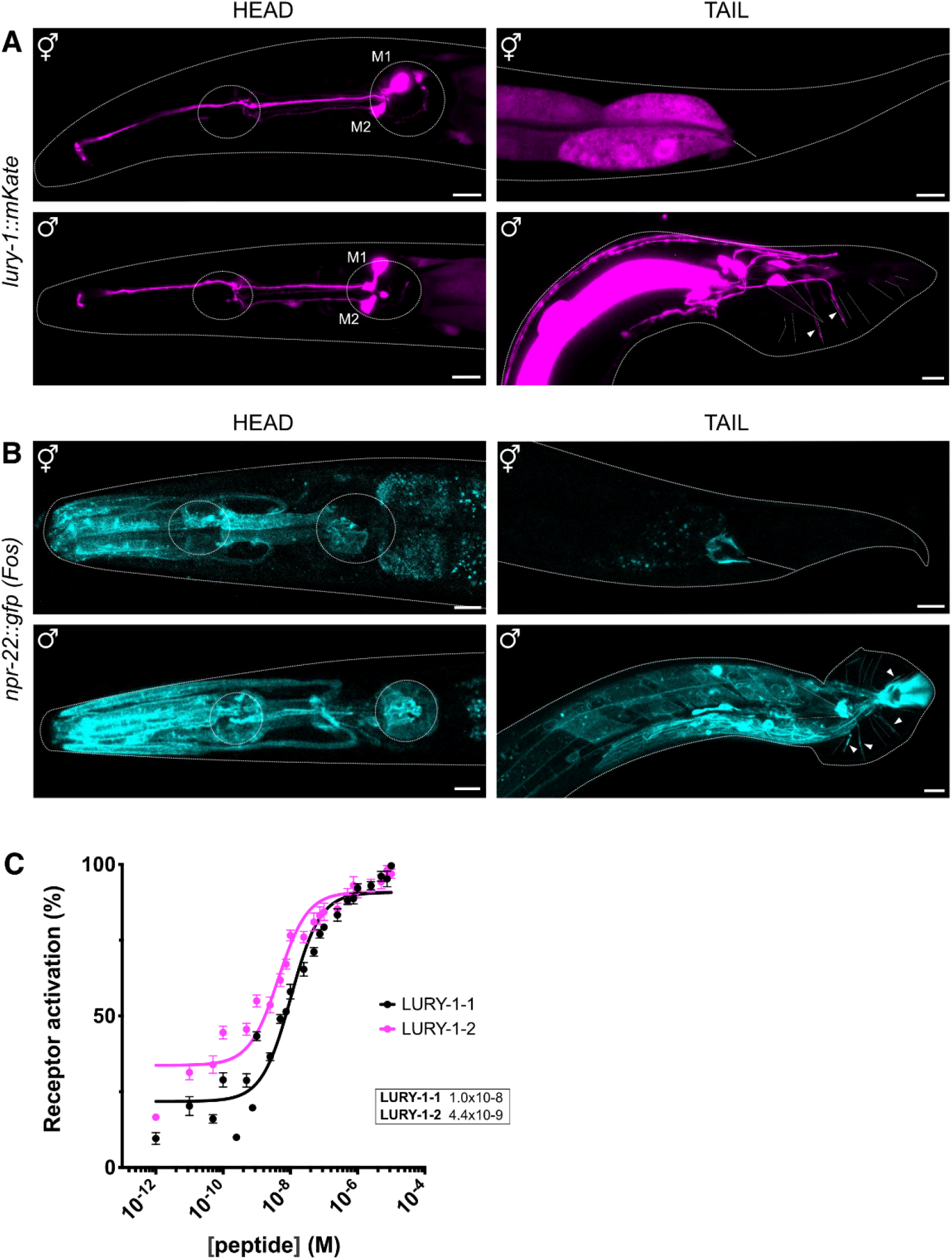
Neuropeptide *lury-1* and its receptor *npr-22* are expressed in different cells in the two sexes of *C. elegans*. **A)** A reporter line expressing the transgene P*lury-1(3.4 kb)∷lury-1 genomic DNA∷SL2-mKate2* shows expression in pharyngeal neurons M1 and M2 (yellow arrows) in both hermaphrodites (top) and males (bottom). In males, there is additional expression in cells in the tail (white arrows). Scale bar = 10 μm. **B)** A reporter line expressing a fosmid containing C-terminal GFP-tagged *npr-22* shows expression in pharynx muscle and cells of the tail in both hermaphrodites (top) and males (bottom). Expression in male copulatory apparatus is also observed. Scale bar = 10 μm. **C)** Dose response curve showing that peptides encoded by the LURY-1 precursor (LURY-1-1 and LURY-1-2) activate the receptor NPR-22 *in vitro.* The corresponding EC50 values (M) for each peptide are as indicated. Line represents non-linear regression fit of a variable slope line using three parameters. n= 6-8 trials.

Given the sex-shared expression of *lury-1* in excitatory pharyngeal neurons known to regulate pharyngeal muscle movements (Pilon & Morck, 2005), we monitored pharyngeal pumping in both hermaphrodites and males (**Figure 2A**). Ohno *et al.* (2017) showed that LURY-1 impacted feeding in hermaphrodites, with *lury-1* over-expression leading to decreased pharyngeal pumping; suggesting that LURY-1 signalling suppresses feeding behaviours (Ohno *et al.*, 2017). In hermaphrodites, we found that mutant animals carrying the *lury-1(gk961835*) allele, a putative null allele that deletes most of the first exon, showed increased pharyngeal pumping (**Figure 2B**). This implies that LURY-1 acts to suppress pharyngeal pumping, consistent with the findings of Ohno *et al.* We did not observe a pharyngeal pumping effect in *npr-22(ok1598)* mutant hermaphrodites (**Figure 2B**), a putative null mutation which encodes a 2.5 kb deletion in the *npr-22* genomic locus (deleting four of six exons). In contrast, male *lury-1* and *npr-22* mutant animals show decreased pharyngeal pumping compared with controls (**Figure 2C**). Male *C. elegans* used in this assay contain a mutation in *him-5* that strongly potentiates the formation of male progeny to ~35% (Meneely *et al*, 2012). Our findings suggest that LURY-1 signalling in males increases pharyngeal pumping, which is opposite to that observed in hermaphrodites. This is curious as pharyngeal neuron expression of *lury-1* is shared between the two sexes, and yet the pharyngeal pumping phenotype of *lury-1* mutants is sexually dimorphic.

**Figure 2:**
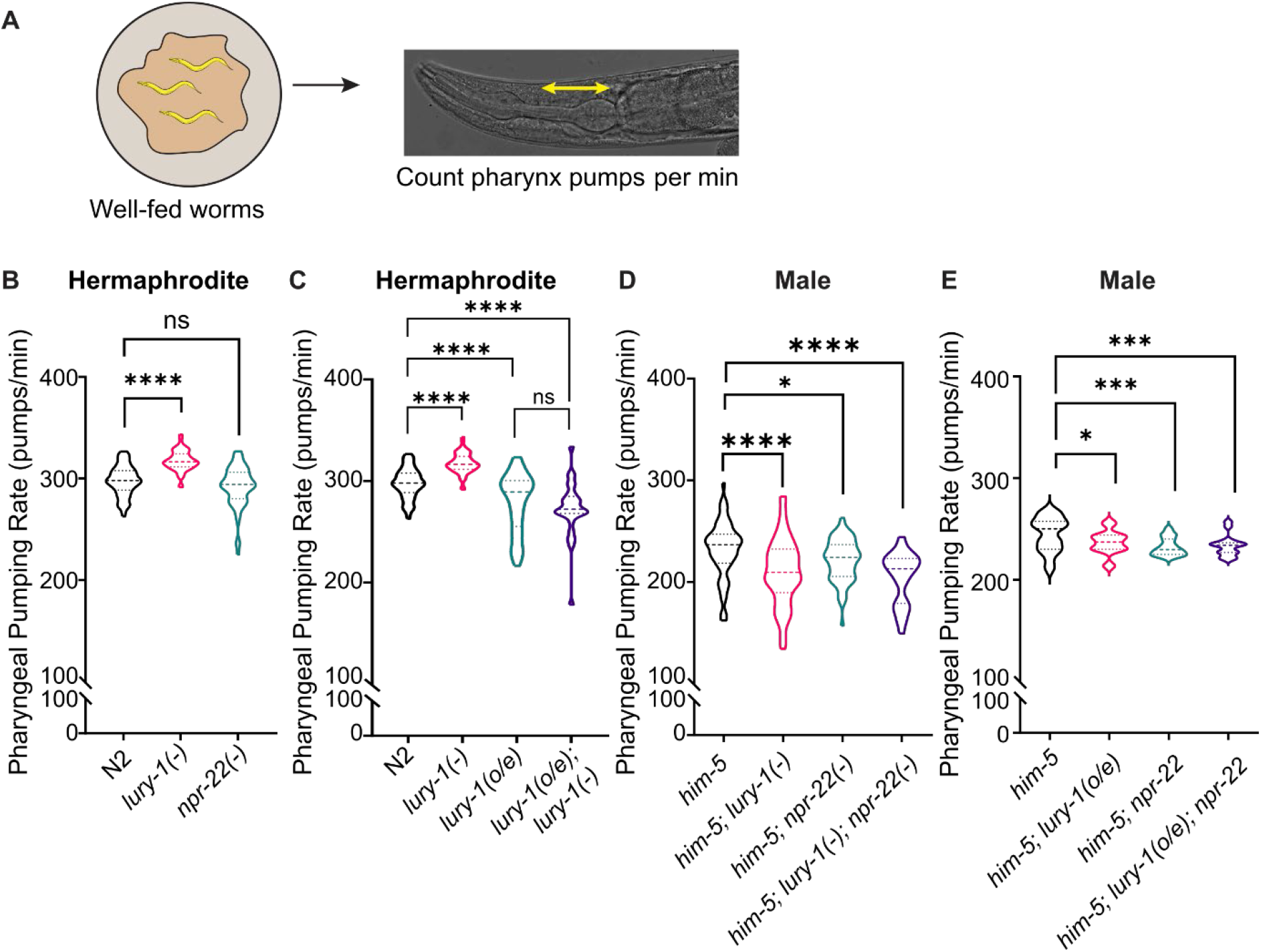
LURY-1 peptides modulate pharyngeal pumping differentially in hermaphrodites and males. **A)** Pharyngeal pumping assays were performed on well-fed animals, with movements of the pharynx grinder counted per minute (pumps/min). **(B-E)** Pharyngeal pumping rate of **(B, D)** *lury-1* and *npr-22* mutant strains and **(C, E)** *lury-1* over-expression (2 kb promoter) transgenic lines containing *lury-1* or *npr-22* mutant alleles. Experiments with male *C. elegans* **(D,E)** were performed in the *him-5* mutant background. n>10 per replicate for >3 biological replicates. Tailed violin plots show median, quartiles and frequency distribution. p-values indicated by ns = not significant. *p<0.05, ***p<0.001, ****p<0.0001 (one-way ANOVA with Fisher’s post-test).

Like the hermaphrodite experiments shown in **Figure 2B**, we also tested pharyngeal pumping rate in male animals over-expressing *lury-1.* In these experiments, we found that *lury-1* over-expression showed reduced pharyngeal pumping in male worms, like that observed in hermaphrodites (**Figure 2D**). These data suggest that, unlike the null mutants, increased gene copy number of *lury-1* has the same effect on pharyngeal pumping in both sexes.

LURY-1 signalling in hermaphrodites modulates egg-laying behaviours (Ohno *et al.*, 2017), indicating a role in reproduction. As we found LURY-1 expression in the tail of male *C. elegans* (**Figure 1B**), and male copulation behaviour is coordinated primarily by neurons of the male tail (Koo *et al*, 2011; Susoy *et al*, 2021), we asked if male reproduction was regulated by LURY-1. To assess male mating efficiency, we picked *fog-2* mutant hermaphrodites, which produce eggs but no sperm – and therefore cannot self-fertilise – onto individual plates with a single male worm (or no male), removing the male after a few hours (see **Methods and Materials**). If no male is present (annotated ‘*no male control*’), or if the male does not mate with the *fog-2(-)* hermaphrodite, there will be no progeny produced (**Figure 3Ai, 3B**). If the male successfully mates with the *fog-2(-)* hermaphrodite, there will be progeny produced through cross-fertilisation (**Figure 3Aii**). We found that *lury-1* and *npr-22* mutant males (both crossed into the *him-5* background) showed more successful mating events than control males (**Figure 3B**). This suggests that LURY-1 signalling suppresses male mating, potentially via NPR-22. We also tested the impact of *lury-1* over-expression on male mating; however, no statistically significant effect of over-expression was observed compared to control males (**Figure EV3**). One possible explanation for this is that transcription of the *lury-1* gene may not be the rate-limiting step in the modulation of male mating, and that post-transcriptional processing or peptide release may be the stage at which regulation of LURY-1 signals are critical in this context.

**Figure 3:**
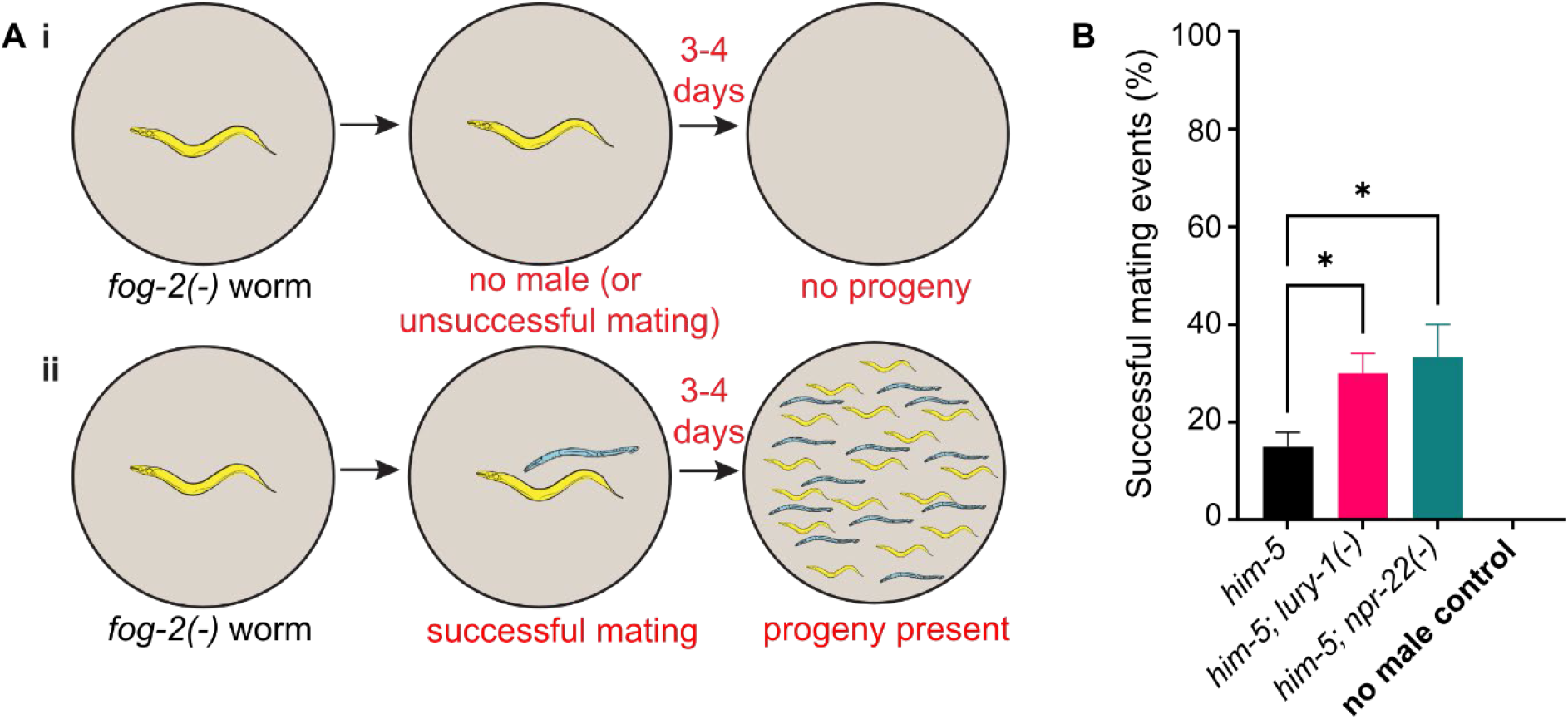
LURY-1 peptides modulate male mating efficiency. **A)** Male mating efficiency assay: (i) *fog-2(-)* produce eggs but no sperm, so will not produce progeny if mating with a male does not occur, or if no male is present. (ii) If mating successfully occurs, then progeny will be observed after several days. **B)** Mating efficiency for *lury-1* and *npr-22* mutant males left to mate with *fog-2(-)* hermaphrodites for 3 hours. Experiments were performed in the *him-5* mutant background. n>10 per genotype, per replicate for >3 biological replicates. Error bars indicate mean ± SEM. p-values indicated by ns = not significant. *p<0.05 (one way ANOVA, Fisher’s post-test).

**Figure EV3:**
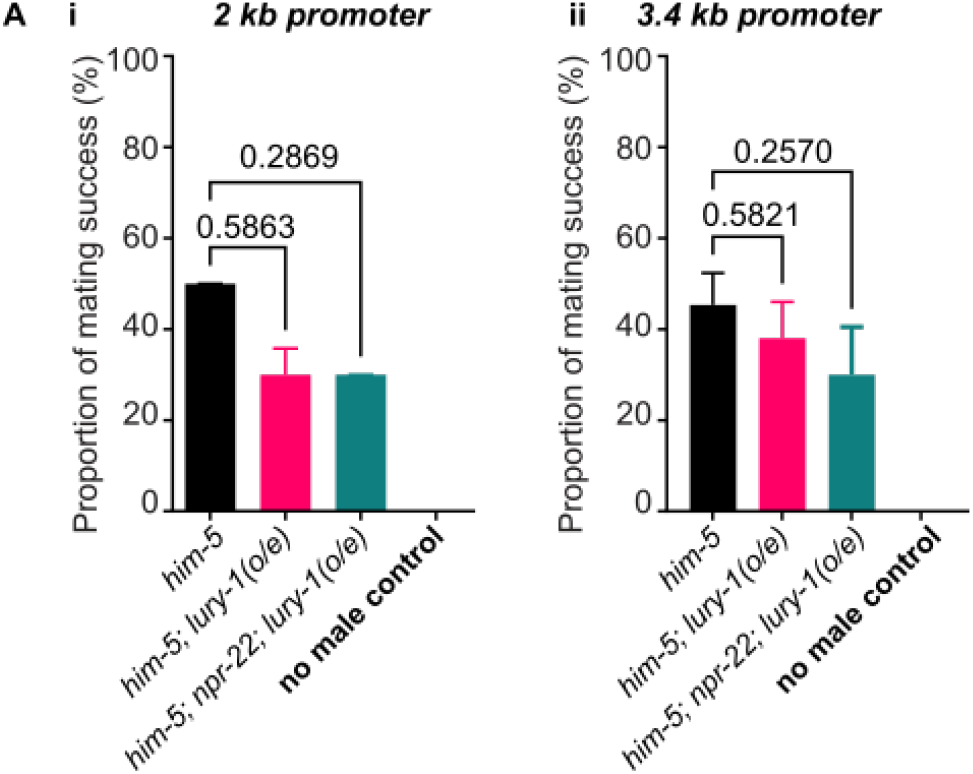
Over-expression of *lury-1* does not affect male mating efficiency. **A)** Mating efficiency for *lury-1* over-expression transgenic lines expressing *lury-1* gDNA using a **(i)** 2 kb promoter (3 hours incubation period with *fog-2(-)* worms) or **(ii)** 3.4 kb promoter (8 hours incubation period with *fog-2(-)* worms). Experiments were performed in the *him-5* mutant background. n>10 per genotype, per replicate for >3 biological replicates. Error bars indicate mean ± SEM. Exact p-values indicated here (one way ANOVA, Fisher’s post-test).

Since *lury-1* mutant males showed a higher mating efficiency compared with controls (**Figure 3B**), we next asked whether LURY-1 affected individual male copulation behaviours. The *C. elegans* mating process is a complex and stereotyped series of behaviours, where the male worm makes contact with the hermaphrodite and scans for the vulva. If it fails to find the vulva, it may turn over the head or tail to scan the other side of the hermaphrodite body (Liu & Sternberg, 1995) (**Figure 4Ai**). Previous reports have carefully characterised the male turning behaviour and categorised it into three (3) types: good, sloppy and missed (Loer & Kenyon, 1993). Good turns occur where contact with the hermaphrodite is never lost, sloppy turns involve a brief loss of contact that is regained, and missed turns are where lost contact is never regained. We recorded mating behaviour of male worms of various genotypes using a worm behavioural tracker (Yemini *et al*, 2013). In general, our data show that most turns are “good”, and that “sloppy” or “missed” turns are relatively infrequent. *lury-1* and *npr-22* mutant animals did not show statistically significant differences in proportion of the different turning categories (**Figure 4B**); however, we did see that *lury-1* over-expression led to a significantly higher proportion of missed turns compared with control wild-type males (**Figure 4C**). This higher proportion of missed turns was not observed when *lury-1* over-expressors were crossed into *lury-1* or *npr-22* putative null mutants, indicating that knocking out the endogenous *lury-1* gene or that of the LURY-1 receptor, ‘rescues’ this phenotype. These data indicate that, in this context, increased transcription of the *lury-1* gene modulates turning during male mating.

**Figure 4:**
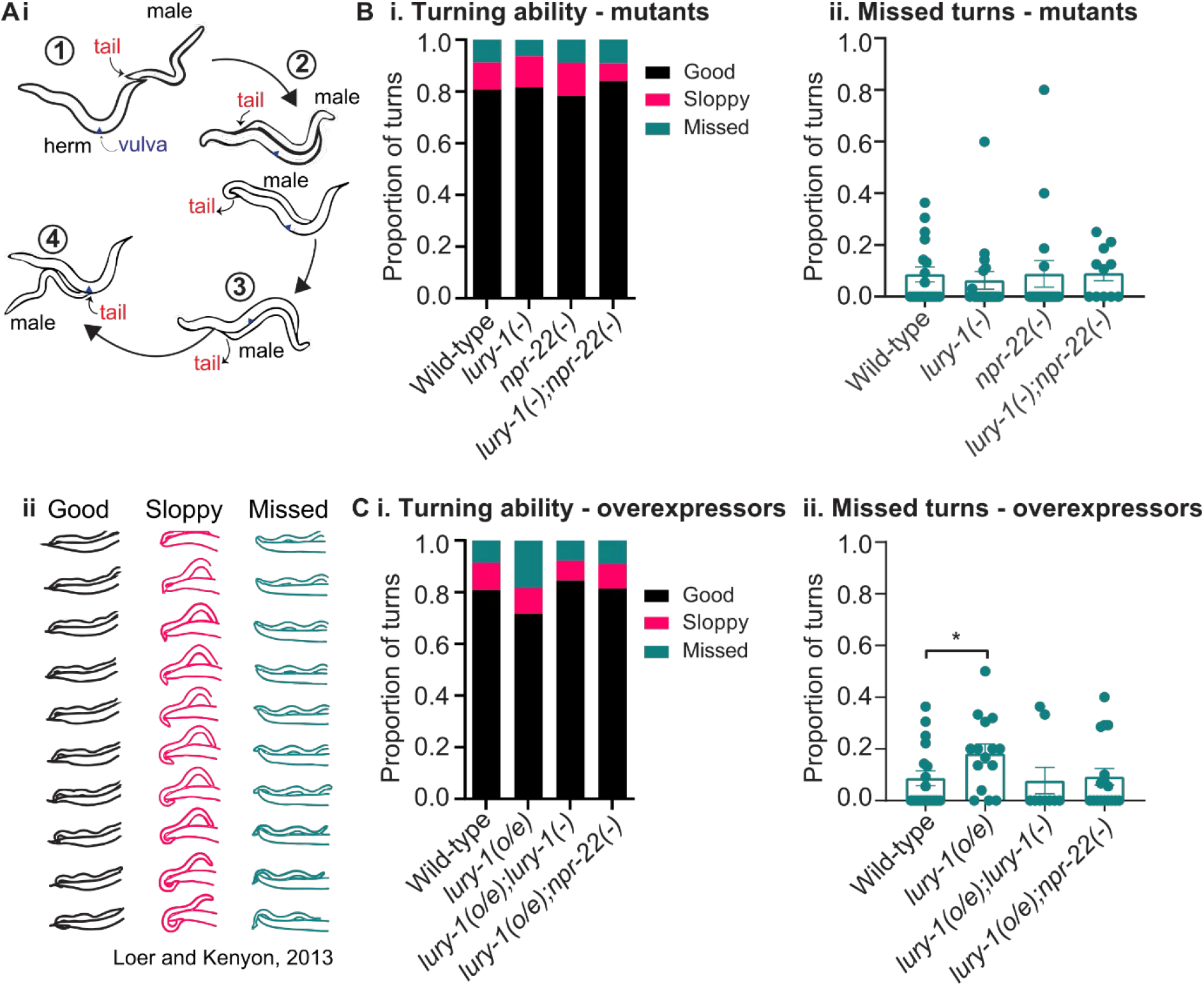
LURY-1 over-expression leads to less efficient turning during male mating. **A) (i)** Male mating behaviour is a complex multi-step process that begins with 1) contacting the hermaphrodite with the male tail and backward movement in search of the vulva; 2) turning and scanning: the male presses its tail against the body of the hermaphrodite and moves backwards until it reaches the vulva. If the male does not detect the vulva, it will make a tight turn and search along the other side; 3) once the vulva is located, the male will prod the slit with his spicules; 4) the male locks to the hermaphrodite and ejaculates his sperm inside. **(ii)** Illustrations of good, sloppy and missed turns, adapted from (Loer & Kenyon, 1993). **B) (i)** Proportion of good, sloppy and missed turns and **(ii)** missed turns as a proportion of all turns as observed in video recordings as observed in video recordings for *lury-1* and *npr-22* mutant males left to mate with an *unc-13* hermaphrodite. Data from B(ii) is extracted from the graph in B(i). From left to right, n = 18, 18, 17, 11. Error bars = mean ± SEM. All comparisons are not statistically significant. **C) (i)** Proportion of good, sloppy and missed turns and **(ii)** missed turns as a proportion of all turns for *lury-1* over-expression transgenic lines crossed with *lury-1* or *npr-22* mutant males left to mate with an *unc-13* hermaphrodite. Data from C(ii) is extracted from the graph in C(i). From left to right, n = 18, 15 , 9, 17. Error bars = mean ± SEM. p-values indicated by *<p=0.05 (one-way ANOVA with Fisher’s post-test).

We next tested whether the nutritional (fed or fasted) status of *lury-1* mutant males affects mating efficiency (**Figure 5A**). Previous reports suggest that fasting male worms increases the likelihood that they will remain on the food patch, in contrast to well-fed males that rapidly leave food to search for mates (Lipton *et al.*, 2004). Using the *fog-2(-)* mating assay (**Figure 3A**), we found that fasting the male worms for 16 hours prior to exposure to *fog-2(-)* hermaphrodites had no statistically significant impact on mating in control or *lury-1* mutant males (**Figure 5Aii**). These data indicate that fed status did not have a strong impact on the modulation of male mating by LURY-1.

**Figure 5.**
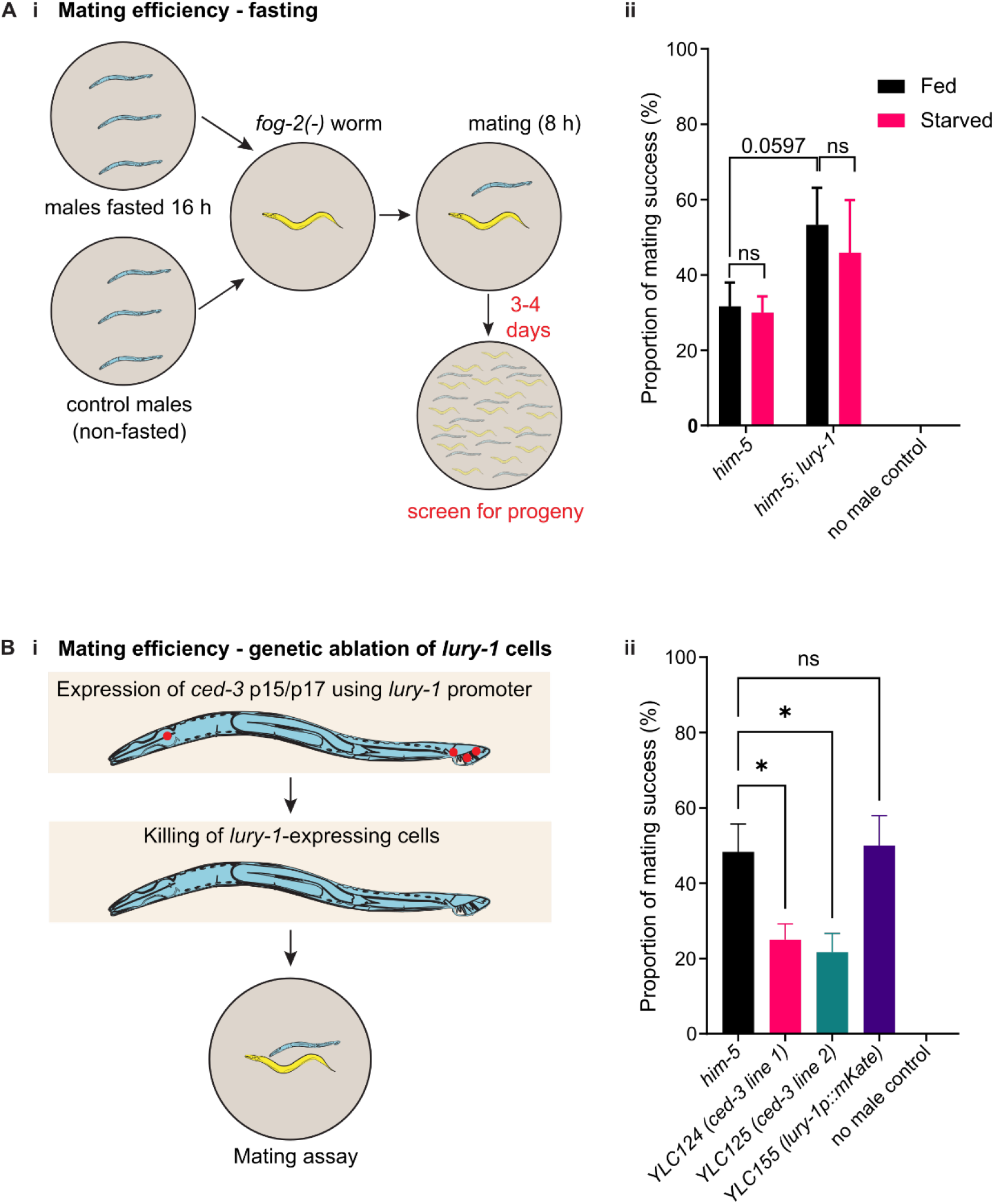
Male mating behaviour as modulated by LURY-1 peptides is not strongly affected by fed state but requires *lury-1*-expressing neurons. **A) (i)** Males are fasted for 16 hours on plates with no food before being moved to plates with *fog-2(-)* hermaphrodites for 8 hours. Control males are not fasted during this period. Males are removed from *fog-2(-)* plates and the presence of progeny on the *fog-2(-)* hermaphrodite plate is assessed after 2-3 days. **(ii)** Mating efficiency for *lury-1* and *npr-22* mutant strains with and without fasting. n=15 per genotype (n=10 for no male control), per replicate for 4 biological replicates. Error bars indicate mean ± SEM. p-values indicated by ns = not significant (two-way ANOVA, Fisher’s post-test). **B) (i)** Caspase *ced-3* p15 and p17 fragments were expressed under the control of the *lury-1* promoter to genetically kill *lury-1-*expressing neurons (Chelur & Chalfie, 2007). **(ii)** Mating efficiency for caspase-killed (*lury-1*p*∷ced-3*, two independent lines) and control (*lury-* 1p∷*mKate2*) males, incubated with *fog-2(-)* hermaphrodites for 8 hours. Experiments were performed in the *him-5* mutant background. n=15 per genotype (n=10 for no male control), per replicate for 4 biological replicates. Error bars indicate mean ± SEM. p-values indicated by ns = not significant. *p<0.05 (one-way ANOVA, Fisher’s post-test).

Expression of *lury-1* was observed in multiple neurons in the male tail (**Figure 1B**). To determine whether these *lury-1*-expressing cells in the male worm regulate mating, we performed genetic ablation of these cells by over-expressing caspase proteins (Chelur & Chalfie, 2007) using the *lury-1* promoter, and testing mating efficiency (**Figure 5Bi**). We found that genetic ablation of *lury-1* cells led to significantly reduced mating efficiency compared with control *him-5* mutant males (**Figure 5Bii**). We also tested a control transgenic line in which *mKate2* was expressed using the *lury-1* promoter and found that this line showed a similar behaviour as control *him-5* males, demonstrating that expression of a non-active transgene using the *lury-1* promoter does not significantly affect male mating (**Figure 5Bii**). Our findings indicate that the cells expressing *lury-1* in the male (pharyngeal neurons, male tail cells) are likely to function in promoting male reproductive behaviours.

In summary, we showed that the *lury-1* neuropeptide precursor is expressed in pharyngeal (feeding) neurons in both *C. elegans* sexes, but with males showing further expression in tail cells required for mating. Interestingly, *lury-1* mutant males show opposing phenotypes in pharyngeal pumping (feeding) behaviours compared with hermaphrodites: LURY-1 in males signals to increase pharyngeal pumping, which is the opposite to its effects in hermaphrodites (Ohno *et al.*, 2017). In addition, LURY-1 influences male mating behaviour, with *lury-1* mutant males showing higher mating efficiency than controls. This modulation of male mating requires *lury-1*-expressing neurons and is not affected by prior starvation. Based on these data, we propose that signalling via LURY-1 peptides may act to regulate both feeding and mating behaviours in a sexually dimorphic manner in *C. elegans*.

Luqin-type neuropeptides are evolutionarily ancient, with the origin of this peptide signalling system predating the divergence of protostomes and deuterostomes (Jekely, 2013; Mirabeau & Joly, 2013). Indeed, luqin peptides have also been implicated in feeding and reproductive behaviours in other invertebrates (reviewed in (Yanez-Guerra & Elphick, 2020)). Our study provides new information on the role of LURY-1 peptides in male *C. elegans*. However, there are several questions that remain to be addressed: Firstly, the effects of *lury-1* over-expression are not as consistent as effects of the null mutation. Although *lury-1* mutant hermaphrodites and *lury-1* over-expressing hermaphrodites show opposing phenotypes, *lury-1* mutant males and *lury-1* over-expressing males show effects in the same direction (both show less pharyngeal pumping) (**Figure 2**). In addition, *lury-1* over-expressing males did not show significant differences in mating efficiency compared to controls in the *fog-2(-)* assay, whereas *lury-1* mutants showed increased mating efficiency. One possible explanation for this is that increasing transcription from the *lury-1* gene may impact the expression of other neuropeptides, with the profile of neuropeptides affected differing between different neuron subtypes (pharyngeal neurons vs male tail neurons). This could then result in different impacts of *lury-1* transgenic over-expression on different behaviours i.e., feeding and reproduction. Moreover, the increased transcription of the peptide precursor may cause excess LURY-1 to be present outside its physiological context, which may lead to other impacts on the biochemistry of these neurons.

Secondly, specifically how LURY-1 release from the neurons of the male tail modulates feeding or mating has not yet been delineated. Recent research using whole-brain imaging and simultaneous behavioural tracking has begun to link individual male tail neurons to specific behaviours during mating (Susoy *et al.*, 2021). Identification of *lury-1* expressing male tail neurons, and characterising their functions, may reveal how LURY-1 release from these neurons and its impacts on mating behaviour are coordinated with release of the same signal from pharyngeal neurons that have not previously been reported to affect male mating.

The sexually dimorphic expression pattern and function of LURY-1 could reflect differences in reproductive behaviours between the two sexes. Hermaphrodites can self-fertilise and therefore do not require mates to reproduce, whereas males only reproduce by mating. Moreover, male *C. elegans* will prioritise mate-seeking over remaining on a source of food, eventually abandoning food to search for mates – a behaviour that hermaphrodites do not exhibit (Lipton *et al.*, 2004; Ryan *et al.*, 2014). LURY-1 in hermaphrodites regulates feeding and egg-laying in a food-dependent manner (Ohno *et al.*, 2017). In males, food-leaving as a strategy for seeking mates involves the coordination of external and internal signals including reproductive and nutritional states, as well as neuromodulatory signalling (Barrios, 2014; Lipton *et al.*, 2004). Previous research has demonstrated the involvement of pigment-dispersing factor (PDF) peptides released from sex-shared (Barrios *et al.*, 2012) and male-specific (Sammut *et al*, 2015) neurons. In addition, nematocin (NTC-1), an oxytocin/vasopressin-related neuropeptide in *C. elegans* (Beets *et al.*, 2012; Garrison *et al.*, 2012), modulates food leaving behaviour and coordination of the multi-step mating behaviour in males (Garrison *et al.*, 2012). In addition to neuropeptides, male food-leaving is also impacted by serotonin and insulin signalling (Lipton *et al.*, 2004), as well as dafachronic acid (DA) signalling through the nuclear hormone receptor DAF-12 (Kleemann *et al*, 2008). Here we have described another peptidergic signal, LURY-1, expressed both in neurons that regulate feeding and those that regulate mating, that may also modulate male copulatory behaviours. Future studies aiming to identify the combined behavioural impact of individual neuromodulator signals on neural circuits in the *C. elegans* male tail will provide a tractable model of how synaptic and “extrasynaptic” signals work together to drive sexually dimorphic behaviours. This may then help to elucidate the general principles through which sexually dimorphic behaviours arise from the combination of neuronal signalling, neuroanatomy, hormonal influence, and experience, relevant to higher organisms.

## Materials and Methods

### Strain Maintenance

Strains were maintained on NGM (nematode growth medium) plates seeded with *E. coli* strain OP50 according to standard experimental procedures (Brenner, 1974). Day 1 adult *C. elegans* were used for all experiments. All experiments were performed at room temperature (22-23 °C) and cultivated at 22 °C.

Mutant strains used include Bristol N2 (wild-type), *fog-2(q71)*, *him-5(e1490), lury-1(gk961835)*, *npr-22(ok1598), him-5(e1490);lury-1(gk961835)* and *him-5(e1490);npr-22(ok1598).* All mutant strains were backcrossed >4 times to N2. A list of transgenic lines used, including full genotype information, is in **Table EV1**. Transgenes were generated through standard microinjection procedures as in (Chew *et al*, 2018a; Chew *et al*, 2018b).

Strain LX2073 (*npr-22∷GFP* fosmid) was a kind gift from the Koelle laboratory (Yale School of Medicine, New Haven, CT). Strains YLC124 and YLC125 (*him-5(e1490);pepEx012[Plury-1(3.4kb)∷ced-3 (p15)∷nz∷unc-54 3’UTR; Plury-1(3.4)∷cz∷ced-3 (p17)∷unc-54 3’UTR; Punc-122∷gfp∷unc-54 3’UTR])* and YLC132 (*him-5;pepEx013[Plury-1(3.4kb)∷mKate2, unc-122∷gfp]*) were generated by SUNY Biotech (Fujian Province, China). Genotypes were confirmed via PCR.

Transgenes were cloned using the Multisite Gateway Three-Fragment cloning system (12537-023, Invitrogen) into pDESTR4R3 II, or using HiFi cloning (NEB #E2621). Promoters for the *lury-1* gene were cloned either 2 kb or 3.4 kb before the ATG. For transgenes including *lury-1* genomic DNA spanning the coding sequence and 3′ UTR, the entire coding region was cloned from the ATG start codon to the TGA stop codon, with an additional 330 bp in the UTR. Lines YL124 and YLC125 were generated by injection of plasmids containing the *ced-3* caspase fragments p15 and p17, expressed under the control of the *lury-1* promoter (3.4 kb). These constructs were generated by replacing the *mec-18* promoter in plasmids TU#806 (Addgene #16080) and TU#807 (Addgene #16081) (Chelur & Chalfie, 2007) with the *lury-1* promoter using Hifi cloning.

### Behavioural Tests

#### Mating Assay

This assay was performed according to the protocol outlined in (Murray et al, 2011) with the following modifications: *fog-2(q71)* hermaphrodites, which do not produce self-progeny, are picked at the fourth larval stage (L4) 16 hours before the assay is conducted. Male *C. elegans* used in this assay contain a mutation in *him-5* that strongly potentiates the formation of male progeny to ~35% (compared with ~0.1% male progeny in wild-type populations). Male worms are also picked at the fourth larval stage (L4) 16 hours before the assay to an OP50-seeded plate with no hermaphrodites present. For the assay, one adult hermaphrodite was transferred onto an individual OP50 seeded plate and one adult male was picked to each plate to begin the mating assay. Males were removed from the plates after 3 or 8 hours, as indicated. The plates were scored as successful or unsuccessful mating based on the presence or absence of progeny on the plates after >1 day. Sample size = n > 10 males per biological replicate for >3 replicates.

For mating assays combined with a period of starvation, assays were adapted as follows: L4 males were picked onto plates with food, or with no food, and left at 22 °C before being placed together with the *fog-2(q71)* hermaphrodite. Males were removed from the plates after 8 hours.

#### Pharyngeal Pumping

The rate of pharyngeal pumping was determined as previously described (Ohno *et al.*, 2017). Adult animals were transferred onto fresh OP50-seeded plates and allowed to acclimatise for 30 min. After 30 min, the number of movements of the grinder in the posterior bulb of the pharynx observed in one minute was counted (pumps/min). Sample size per genotype n > 10 per biological replicate for 3 replicates.

#### Turn scoring assay

Preparation of plates and recording procedures were performed as in (Yan *et al*, 2017; Yemini *et al.*, 2013) with the following changes: Video recordings were performed using a single *unc-13* hermaphrodite placed on food on a 3 cm NGM plate. *unc-13* hermaphrodites were used for ease of tracking – these animals have severely uncoordinated locomotion and do not move across the plate, allowing us to track only the freely-moving (male) worm. After ~5 minutes, a male worm was picked onto this plate. Plates were placed on the tracker and recordings were started 15 minutes later. Each recording was 30 minutes. Genotypes were blinded during recording and analysis. Scoring the number of Good, Sloppy and Missed turns was performed according to (Loer & Kenyon, 1993), with the following conditions: male was in contact with hermaphrodite for >5 minutes for the duration of the recording, 1-3 contacts were recorded per animal. n > 10 per genotype.

### In vitro *GPCR activation assays*

Cell-based activation assays were performed as described (Beets *et al.*, 2012). NPR-22 cDNA was cloned into the pcDNA3.1 TOPO expression vector (Thermo Fisher Scientific). Receptor activation was studied in Chinese hamster ovary cells (CHO) stably expressing apo-aequorin (mtAEQ) targeted to the mitochondria as well as the human Gα16 subunit. The CHO-K1 cell line (PerkinElmer, ES-000-A24) was used for receptor activation assays. CHO/mtAEQ/Gα16 cells were transiently transfected with the NPR-22 cDNA construct or the empty pcDNA3.1 vector using the Lipofectamine transfection reagent (Thermo Fisher Scientific). Cells expressing the receptor were shifted to 28°C 1 day later, and collected 2 days post-transfection in BSA medium (DMEM/HAM’s F12 with 15 mM HEPES, without phenol red, 0.1% BSA) loaded with 5 μM coelenterazine h (Thermo Fisher Scientific) for 4 h to reconstitute the holo-enzyme aequorin. Cells (25,000 cells/well) were exposed to synthetic peptides in BSA medium, and aequorin bioluminescence was recorded for 30 s on a MicroBeta LumiJet luminometer (PerkinElmer, Waltham Massachusetts) in quadruplicate. For dose-response evaluations, after 30 s of ligand-stimulated calcium measurements, Triton X-100 (0.1%) was added to the well to obtain a measure of the maximum cell Ca^2+^ response. BSA medium without peptides was used as a negative control and 1 μM ATP was used to check the functional response of the cells. Cells transfected with the empty vector were used as a negative control (not shown). EC50 values were calculated from dose-response curves, constructed using a nonlinear regression analysis, with sigmoidal dose-response equation (Prism 9.0).

### Confocal microscopy

Confocal images were obtained using a Leica SP8 confocal microscope (University of Wollongong) and Zeiss LSM880 (Flinders University). Z-stacks were analysed using Fiji (ImageJ).

### Statistical Analyses

Statistical analysis for all experiments was performed using GraphPad Prism 8.0. In general, where two groups were compared, an unpaired t-test was used. Where multiple groups tested with a single condition were compared, a one-way ANOVA with Fisher’s post-test was used. Where multiple groups tested with multiple conditions were compared, a two-way ANOVA with Fisher’s post-test was used. The alpha value for all analyses is set at 0.05.

## Acknowledgments

Many thanks go to the members of the Chew and Schafer labs for helpful discussions in the preparation of this manuscript. We specifically thank Aelon Rahmani for proofreading the manuscript. We gratefully acknowledge the *Caenorhabditis* Genetics Centre, which is supported by the National Institutes of Health (P40 OD010440), for providing some of the strains used in this study. We thank Prof Michael Koelle (Yale School of Medicine) for providing strain LX2073.

## Author contributions

Conceptualisation/Methodology/Project administration/Supervision – **IB, IH, WRS, YLC**. Data curation/Formal analysis/Investigation – **MJG, IH, DS, PJ, IB, YLC.** Funding acquisition – **IB, WRS, YLC.** Visualisation/Writing – original draft/Writing – review & editing – **IB, IH, WRS, YLC**

## Data Availability Section

This study includes no data deposited in external repositories. Reagents including *C. elegans* strains and expression plasmids are available from the corresponding author upon request.

---

**Table EV1:** List of strains used in this study. For transgenic lines, the number following each construct in brackets indicates the ng/μL of plasmid injected e.g. (50) means 50 ng/μL of the construct was included in the injection mix.

| Strain number | Genotype |
| --- | --- |
| N2 | Wild-type |
| AQ4214 | *fog-2(q71)* |
| AQ4266 | *him-5(e1490)V bc4x* |
| YLC022 | *him-5(e1490) V; lury-1(gk961835)* |
| AQ4287 | *him-5(e1490)V; npr-22(ok1598) bc 5x* |
| AQ4269 | *him-5(e1490)V bc4x; npr-22(ok1598) bc 5x; lury-1(gk961835)* |
| AQ4291 | *him-5(e1490)V; lJEx1001[Plury-1(2kb)::lury-1::SL2-mKate2(25);unc-122::gfp(50)]* |
| AQ4215 | *lury-1(gk961835);lJEx1001[Plury-1(2kb)::lury-1::SL2-mKate2(25);unc-122::gfp(50)]* |
| AQ4270 | *him-5(e1490)V bc4x; npr-22(ok1598) bc 5x; lJEx1001[Plury-1(2kb)::lury-1::SL2-mKate2(25);unc-122::gfp(50)]* |
| AQ4290 | *him-5(e1490)V; bc4x; lury-1(gk961835) backcrossed 6x; lJEx1001[Plury-1(2kb)::lury-1::SL2-mKate2(25);unc-122::gfp(50)]* |
| YLC103 | *him-5(e1490); pepEx009[Plury-1(3.4kb)::lury-1 gDNA + UTR::SL2-mKate2(pYLC016)(50); unc-122::gfp (60)]- line 1* |
| YLC104 | *him-5(e1490); npr-22(ok1598); pepEx009[Plury-1(3.4kb)::lury-1 gDNA + UTR::SL2-mKate2(pYLC016)(50); unc-122::gfp (60)]- line 1* |
| YLC105 | *him-5(e1490); lury-1(gk961835); pepEx009[Plury-1(3.4kb)::lury-1 gDNA + UTR::SL2-mKate2(pYLC016)(50); unc-122::gfp (60)]- line 1* |
| YLC124 | *him-5(e1490); pepEx011[Plury-1(3.4kb)::ced-3(p15)::nz::unc-54 3'UTR; Plury-1(3.4)::cz::ced-3(p17)::unc-54 3'UTR; Punc-122::gfp::unc-54 3'UTR] line 1* |
| YLC125 | *him-5(e1490);pepEx012[Plury-1(3.4kb)::ced-3(p15)::nz::unc-54 3'UTR; Plury-1(3.4)::cz::ced-3(p17)::unc-54 3'UTR; Punc-122::gfp::unc-54 3'UTR] line 2* |
| YLC132 | *him-5;pepEx013[Plury-1(3.4kb)::mKate2, unc-122::gfp]* |
| LX2073 | *npr-22::GFP fosmid* |

## References

Asahina K, Watanabe K, Duistermars BJ, Hoopfer E, Gonzalez CR, Eyjolfsdottir EA, Perona P, Anderson DJ (2014) Tachykinin-expressing neurons control male-specific aggressive arousal in Drosophila. Cell 156: 221–235

Bargmann CI (2012) Beyond the connectome: how neuromodulators shape neural circuits. Bioessays 34: 458–465

Barr MM, Garcia LR, Portman DS (2018) Sexual Dimorphism and Sex Differences in Caenorhabditis elegans Neuronal Development and Behavior. Genetics 208: 909–935

Barrios A (2014) Exploratory decisions of the Caenorhabditis elegans male: a conflict of two drives. Semin Cell Dev Biol 33: 10–17

Barrios A, Ghosh R, Fang C, Emmons SW, Barr MM (2012) PDF-1 neuropeptide signaling modulates a neural circuit for mate-searching behavior in C. elegans. Nat Neurosci 15: 1675–1682

Bayer EA, Stecky RC, Neal L, Katsamba PS, Ahlsen G, Balaji V, Hoppe T, Shapiro L, Oren-Suissa M, Hobert O (2020) Ubiquitin-dependent regulation of a conserved DMRT protein controls sexually dimorphic synaptic connectivity and behavior. Elife 9

Beets I, Janssen T, Meelkop E, Temmerman L, Suetens N, Rademakers S, Jansen G, Schoofs L (2012) Vasopressin/oxytocin-related signaling regulates gustatory associative learning in C. elegans. Science 338: 543–545

Bentley B, Branicky R, Barnes CL, Chew YL, Yemini E, Bullmore ET, Vertes PE, Schafer WR (2016) The Multilayer Connectome of Caenorhabditis elegans. PLoS Comput Biol 12: e1005283

Brenner S (1974) The genetics of Caenorhabditis elegans. Genetics 77: 71–94

Chelur DS, Chalfie M (2007) Targeted cell killing by reconstituted caspases. Proc Natl Acad Sci U S A 104: 2283–2288

Chew YL, Grundy LJ, Brown AEX, Beets I, Schafer WR (2018a) Neuropeptides encoded by *nlp-49* modulate locomotion, arousal and egg-laying behaviours in *Caenorhabditis elegans* via the receptor SEB-3. Philos Trans R Soc Lond B Biol Sci 373

Chew YL, Tanizawa Y, Cho Y, Zhao B, Yu AJ, Ardiel EL, Rabinowitch I, Bai J, Rankin CH, Lu H et al (2018b) An Afferent Neuropeptide System Transmits Mechanosensory Signals Triggering Sensitization and Arousal in C. elegans. Neuron 99: 1233–1246 e1236

Clayton JA (2016) Sex influences in neurological disorders: case studies and perspectives. Dialogues Clin Neurosci 18: 357–360

Cook SJ, Jarrell TA, Brittin CA, Wang Y, Bloniarz AE, Yakovlev MA, Nguyen KCQ, Tang LT, Bayer EA, Duerr JS et al (2019) Whole-animal connectomes of both Caenorhabditis elegans sexes. Nature 571: 63–71

Fairbairn DJ (1997) Allometry for sexual size dimorphism: Pattern and process in the coevolution of body size in males and females. Annu Rev Ecol Syst 28: 659–687

Garrison JL, Macosko EZ, Bernstein S, Pokala N, Albrecht DR, Bargmann CI (2012) Oxytocin/vasopressin-related peptides have an ancient role in reproductive behavior. Science 338: 540–543

Jekely G (2013) Global view of the evolution and diversity of metazoan neuropeptide signaling. Proc Natl Acad Sci U S A 110: 8702–8707

Kleemann G, Jia L, Emmons SW (2008) Regulation of Caenorhabditis elegans male mate searching behavior by the nuclear receptor DAF-12. Genetics 180: 2111–2122

Koo PK, Bian X, Sherlekar AL, Bunkers MR, Lints R (2011) The robustness of Caenorhabditis elegans male mating behavior depends on the distributed properties of ray sensory neurons and their output through core and male-specific targets. J Neurosci 31: 7497–7510

Lipton J, Kleemann G, Ghosh R, Lints R, Emmons SW (2004) Mate searching in Caenorhabditis elegans: a genetic model for sex drive in a simple invertebrate. J Neurosci 24: 7427–7434

Liu KS, Sternberg PW (1995) Sensory regulation of male mating behavior in Caenorhabditis elegans. Neuron 14: 79–89

Liu T, Kim K, Li C, Barr MM (2007) FMRFamide-like neuropeptides and mechanosensory touch receptor neurons regulate male sexual turning behavior in Caenorhabditis elegans. J Neurosci 27: 7174–7182

Loer CM, Kenyon CJ (1993) Serotonin-deficient mutants and male mating behavior in the nematode Caenorhabditis elegans. J Neurosci 13: 5407–5417

McCarthy MM, Arnold AP (2011) Reframing sexual differentiation of the brain. Nat Neurosci 14: 677–683

Meneely PM, McGovern OL, Heinis FI, Yanowitz JL (2012) Crossover distribution and frequency are regulated by him-5 in Caenorhabditis elegans. Genetics 190: 1251–1266

Mirabeau O, Joly JS (2013) Molecular evolution of peptidergic signaling systems in bilaterians. Proc Natl Acad Sci U S A 110: E2028–2037

Ohno H, Yoshida M, Sato T, Kato J, Miyazato M, Kojima M, Ida T, Iino Y (2017) Luqin-like RYamide peptides regulate food-evoked responses in C. elegans. Elife 6

Palamiuc L, Noble T, Witham E, Ratanpal H, Vaughan M, Srinivasan S (2017) A tachykinin-like neuroendocrine signalling axis couples central serotonin action and nutrient sensing with peripheral lipid metabolism. Nat Commun 8: 14237

Pilon M, Morck C (2005) Development of Caenorhabditis elegans pharynx, with emphasis on its nervous system. Acta Pharmacol Sin 26: 396–404

Portman DS (2017) Sexual modulation of sex-shared neurons and circuits in Caenorhabditis elegans. J Neurosci Res 95: 527–538

Reilly DK, McGlame EJ, Vandewyer E, Robidoux AN, Muirhead CS, Northcott HT, Joyce W, Alkema MJ, Gegear RJ, Beets I et al (2021) Distinct neuropeptide-receptor modules regulate a sex-specific behavioral response to a pheromone. Commun Biol 4: 1018

Ryan DA, Miller RM, Lee K, Neal SJ, Fagan KA, Sengupta P, Portman DS (2014) Sex, age, and hunger regulate behavioral prioritization through dynamic modulation of chemoreceptor expression. Curr Biol 24: 2509–2517

Sammut M, Cook SJ, Nguyen KCQ, Felton T, Hall DH, Emmons SW, Poole RJ, Barrios A (2015) Glia-derived neurons are required for sex-specific learning in C. elegans. Nature 526: 385–390

Susoy V, Hung W, Witvliet D, Whitener JE, Wu M, Park CF, Graham BJ, Zhen M, Venkatachalam V, Samuel ADT (2021) Natural sensory context drives diverse brain-wide activity during C. elegans mating. Cell 184: 5122–5137 e5117

Tabatadze N, Huang G, May RM, Jain A, Woolley CS (2015) Sex Differences in Molecular Signaling at Inhibitory Synapses in the Hippocampus. J Neurosci 35: 11252–11265

Turek M, Besseling J, Spies JP, Konig S, Bringmann H (2016) Sleep-active neuron specification and sleep induction require FLP-11 neuropeptides to systemically induce sleep. Elife 5

Walker DS, Chew YL, Schafer WR, 2017. Genetics of Behavior in C. elegans, in: Byrne J.H. (Ed.) The Oxford Handbook of Invertebrate Neurobiology. Oxford University Press.

Wexler LR, Miller RM, Portman DS (2020) C. elegans Males Integrate Food Signals and Biological Sex to Modulate State-Dependent Chemosensation and Behavioral Prioritization. Curr Biol 30: 2695–2706 e2694

White JG, Southgate E, Thomson JN, Brenner S (1986) The structure of the nervous system of the nematode Caenorhabditis elegans. Philos Trans R Soc Lond B Biol Sci 314: 1–340

Witvliet D, Mulcahy B, Mitchell JK, Meirovitch Y, Berger DR, Wu Y, Liu Y, Koh WX, Parvathala R, Holmyard D et al (2021) Connectomes across development reveal principles of brain maturation. Nature 596: 257–261

Yan G, Vertes PE, Towlson EK, Chew YL, Walker DS, Schafer WR, Barabasi AL (2017) Network control principles predict neuron function in the Caenorhabditis elegans connectome. Nature 550: 519–523

Yanez-Guerra LA, Elphick MR (2020) Evolution and Comparative Physiology of Luqin-Type Neuropeptide Signaling. Front Neurosci 14: 130

Yemini E, Jucikas T, Grundy LJ, Brown AE, Schafer WR (2013) A database of Caenorhabditis elegans behavioral phenotypes. Nat Methods 10: 877–879

